# The architecture of cell differentiation in choanoflagellates and sponge choanocytes

**DOI:** 10.1101/452185

**Authors:** Davis Laundon, Ben Larson, Kent McDonald, Nicole King, Pawel Burkhardt

## Abstract

Collar cells are ancient animal cell types which are conserved across the animal kingdom [1] and their closest relatives, the choanoflagellates [2]. However, little is known about their ancestry, their subcellular architecture, or how they differentiate. The choanoflagellate *Salpingoeca rosetta* [3] expresses genes necessary for animal multicellularity and development [4] and can alternate between unicellular and multicellular states [3,5], making it a powerful model to investigate the origin of animal multicellularity and mechanisms underlying cell differentiation [6,7]. To compare the subcellular architecture of solitary collar cells in *S. rosetta* with that of multicellular “rosettes” and collar cells in sponges, we reconstructed entire cells in 3D through transmission electron microscopy on serial ultrathin sections. Structural analysis of our 3D reconstructions revealed important differences between single and colonial choanoflagellate cells, with colonial cells exhibiting a more amoeboid morphology consistent with relatively high levels of macropinocytotic activity. Comparison of multiple reconstructed rosette colonies highlighted the variable nature of cell sizes, cell-cell contact networks and colony arrangement. Importantly, we uncovered the presence of elongated cells in some rosette colonies that likely represent a distinct and differentiated cell type. Intercellular bridges within choanoflagellate colonies displayed a variety of morphologies and connected some, but not all, neighbouring cells. Reconstruction of sponge choanocytes revealed both ultrastructural commonalities and differences in comparison to choanoflagellates. Choanocytes and colonial choanoflagellates are typified by high amoeboid cell activity. In both, the number of microvilli and volumetric proportion of the Golgi apparatus are comparable, whereas choanocytes devote less of their cell volume to the nucleus and mitochondria than choanoflagellates and more of their volume to food vacuoles. Together, our comparative reconstructions uncover the architecture of cell differentiation in choanoflagellates and sponge choanocytes and constitute an important step in reconstructing the cell biology of the last common ancestor of the animal kingdom.

## RESULTS AND DISCUSSION

### Three-dimensional cellular architecture of choanoflagellates

Collar cells were likely one of the first animal cell types [1,8,9] and persist in most animal phyla (Figure 1A). Therefore, characterising the microanatomy of choanoflagellates and sponge choanocytes has important implications for the origin and evolution of animal cell types. To fully characterise and reconstruct both single and colonial *S. rosetta* cells, we used high-pressure freezing and 3D serial ultrathin TEM sectioning (3D ssTEM), in addition to fluorescent microscopy. Three randomly selected single cells and three randomly selected colonial cells from a single colony were chosen for the reconstruction of entire choanoflagellate cells and subcellular structures (Figures 1 and S1-2, Videos S1-6). Both single and colonial *S. rosetta* cells exhibited a prominent, central nucleus enveloped by a mitochondrial reticulum and basal food vacuoles – as well as intracellular glycogen reserves - consistent with the coarse choanoflagellate cellular architecture reported in previous studies [10,11] (reviewed in [7,12]) (Figures 1 and S1-2, Videos S1-6). However, with the increased resolution of electron microscopy we detected three morphologically distinct populations of intracellular vesicles with distinct subcellular localizations (Figure 1G and S1): 1) Large vesicles (extremely electron-lucent, 226 ± 53 nm in diameter), 2) Golgi-associated vesicles (electron-dense inclusions, 50 ± 10 nm in diameter), and 3) Apical vesicles (electron-lucent, 103 ± 21 nm in diameter). Extracellular vesicles were also observed associated with two of the single cells (electron-lucent, 173 ± 36 nm in diameter) and appeared to bud from the microvillar membrane (Fig S1L). Choanoflagellate cells subjected to fluorescent labelling were congruent with 3D ssTEM reconstructions in terms of organelle localization (Figure 1B-C), providing evidence that the 3D models presented herein are biologically representative.

**Figure 1.**
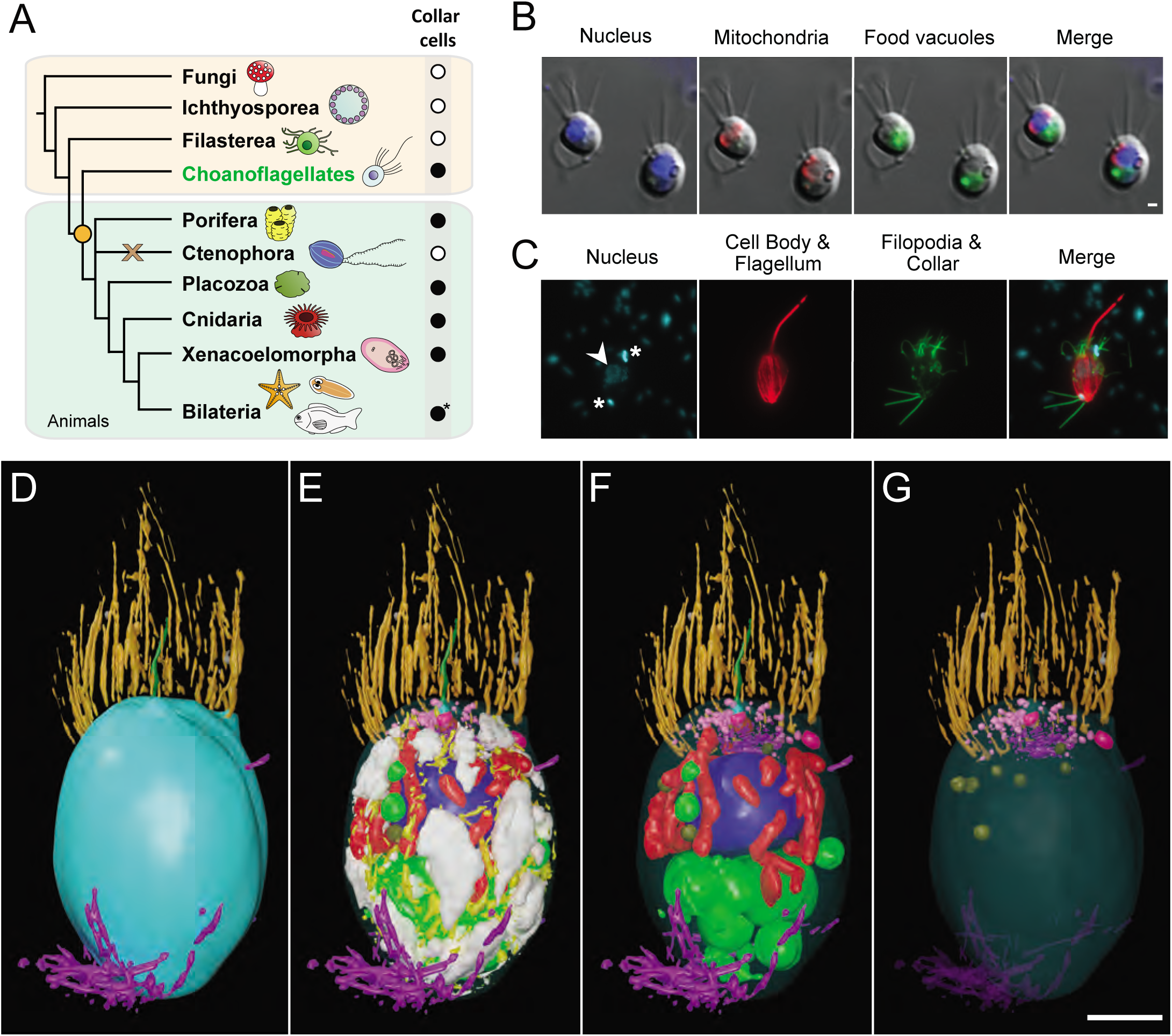
Three-dimensional cellular architecture of choanoflagellates and collar cells across the Choanozoa. (A) Phylogenetic distribution of collar cells across the Choanozoa (Choanoflagellata + Animals [1][44]) showing the presence (black circle), absence (white circle) and putative losses (brown cross) of collar cells across lineages. The origin of collar cells is marked by the orange circle. Adapted from [1]. *Some lineages within the Bilateria have secondarily lost collar cells. (B-C) Characterisation of major organelles in *S. rosetta* labelled with fluorescent vital dyes (B) and by immunofluorescence (C). Arrowhead indicates nucleus of choanoflagellates cell, asterisks indicate the stained nucleoids of engulfed prey bacteria. Scalebar = 1 μm. (D-G) 3D ssTEM reconstruction of a single *S. rosetta* cell (S3) exterior (D). The plasma membrane was made transparent (E) and glycogen and ER were removed to allow better visualisation of subcellular structures (F) and vesicle populations (G). Shown are apical vesicles (pink), food vacuoles (green), endocytotic vacuoles (fuschia), endoplasmic reticulum (yellow), extracellular vesicles (grey), filopodia (external – purple), flagellar basal body (light blue), flagellum (dark green), glycogen storage (white), Golgi apparatus and vesicles (purple), intercellular bridges (external – yellow; septa - red), large vesicles (brown), microvillar collar (light orange), mitochondria (red), non-flagellar basal body (dark orange) and nuclei (dark blue). Scale bar = ∼1 μm (depending on position of structure along the z-axis).

### Ultrastructural commonalities and differences between single and colonial choanoflagellate cells

Our 3D ssTEM reconstructions allowed for detailed volumetric and numerical comparisons among single and colonial *S. rosetta* cells (Figures 2 and S2, Table S1 and S2). Overall, the general deposition of organelles was unchanged in both cell types (Figures 2A, B and S2A-C). In addition, single and colonial cells devote a similar proportion of cell volume to most of their major organelles (nucleus: single cells 12.92 ± 0.58% vs colonial cells 11.56 ± 0.27%; nucleolus: 1.85 ± 0.33% vs 2.2 ± 0.22%; mitochondria: 5.08 ± 1.14% vs 6.63 ± 0.42%; food vacuoles: 9.22 ± 2.75% vs 6.85 ± 0.87% and glycogen storage: 8.71 ± 2.36% vs 7.50 ± 1.12%) (Figures 2 and S2, Table S1 and S2).

**Figure 2.**
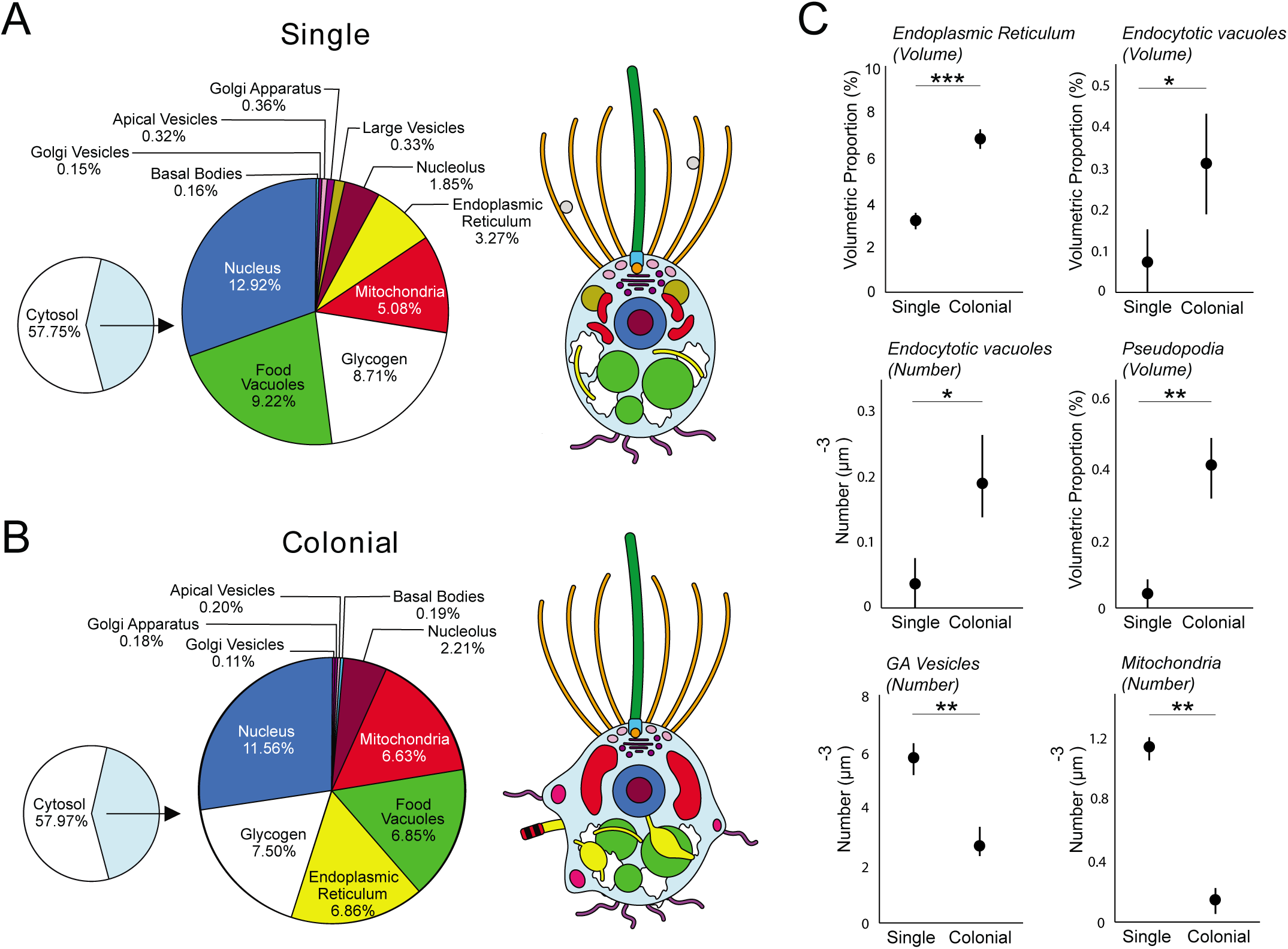
3D ssTEM reconstructions allow for volumetric and numerical comparison of high-resolution single and colonial *S. rosetta* cells. Shown are the mean volumetric breakdowns of three single (A) and three colonial (B) *S. rosetta* cells (left) and a generalised diagram of cell-type ultrastructure (right). Colours are as in Figure 1. (C) Volumetric (%) (±SEM) (endoplasmic reticulum and endocytotic vacuoles) and numerical (μm^-3^) (±SEM) (endocytotic vacuoles, pseudopodia, Golgiassociated vesicles and mitochondria) differences were found between single and colonial (*n* = 3) *S. rosetta* cells. * *p* < 0.05, ** *p* < 0.01, *** *p* < 0.001.

We also uncovered some ultrastructural differences between single and colonial cells (Figure 2C). Colonial cells devoted a higher proportion of cell volume to endoplasmic reticulum (ER) (single: 3.27 ± 0.35% vs colonial: 6.86 ± 0.39%). This contrast was coupled to a differential ER morphology across cell types. The ER of colonial cells frequently displayed wide, flat sheets (Figure 3E), which were not observed in the reconstructed single cells. Single cells exhibited a higher number of Golgi-associated vesicles (single: 166.3 ± 32.7 vs colonial: 72.3 ± 26.5) and individual mitochondria than colonial cells (single: 25.3 ± 5.8 vs colonial: 4.3 ± 4.2) (Figure 2C, Table S2), despite lacking volumetric differences between cell types. The ultrastructural differences in ER, Golgi-associated vesicles and mitochondria suggest differences in endomembrane trafficking and energetic physiology between single and colonial cells. ER and mitochondrial morphology change dynamically, and stark changes have been observed in other eukaryotic cells due to changes in cell cycle [13] and cytoskeletal activity [14,15]. Mitochondria and the ER too show an intimate association [16], and the contrast in the number of individual mitochondria in different cell types was particularly striking (Table S2).

**Figure 3.**
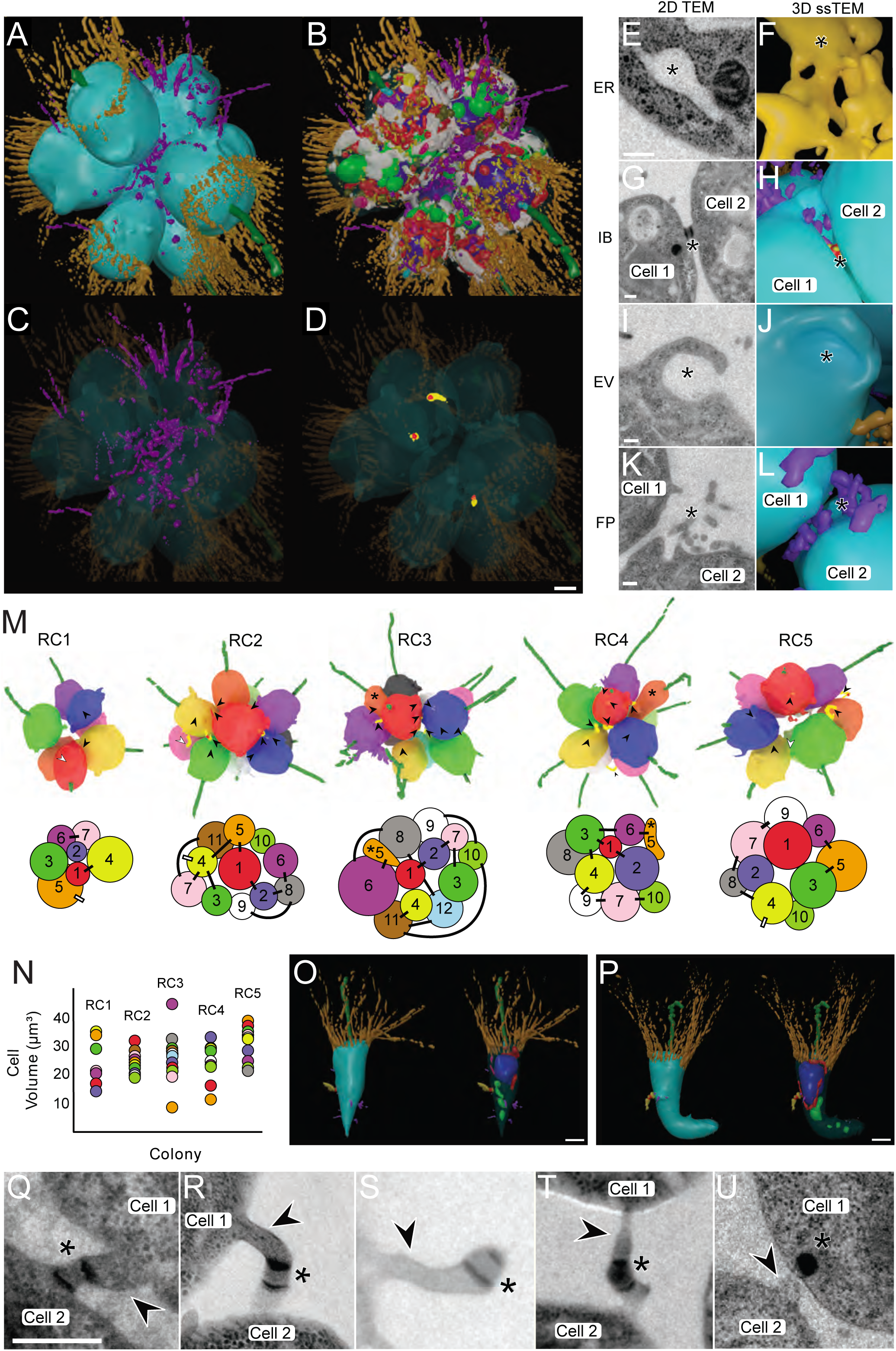
Reconstructions of complete choanoflagellate rosette colonies places colonial cells into context, unveils ultrastructural features involved in rosette formation and a novel cell type. (A-D) 3D ssTEM reconstruction of a complete rosette colony 1 (RC1). The plasma membrane was made transparent (B) to allow better visualisation of subcellular structures. Highlighted are contacting filopodia (C) and intercellular bridges (D). Cellular structures coloured as in Figure 1. Scalebar = ∼1 μm. (E-L) 2D TEM and 3D ssTEM reconstructions of structures (*) differentially exhibited by colonial cells or involved in colony formation. Shown are the endoplasmic reticulum (ER) (E&F), intercellular bridges (IB)(G&H), endocytotic vacuoles (EV) (I&J) and filopodia (FP) (K&L). Scalebars = 200 nm. (M-P) Reconstruction of multiple *S. rosetta* colonies shows no strong pattern of volumetric distribution and bridge networks, but reveal the presence of highly derived cell morphologies. (M) 3D ssTEM reconstructions of five complete rosettes (RC1-5) coloured by cell number (above) and 2D projections of bridge connections in 3D ssTEM reconstructions of rosette colonies (below). Disconnected intercellular bridges marked by white arrowheads and lines. Asterisks mark the presence of highly derived cell morphologies in RC3 and RC4. Cells in rosette colonies are numbered in order of their appearance along the z-axis. (N) Volumetric distribution of mean cell volumes (RC1-5) in rosette colonies reveals no apparent pattern of cell distribution across the z-axis. (O, P). Two highly derived cell types, the ‘carrot cell’ (O) from RC3 and the ‘chili cell’ (P) from RC4 were identified in rosette colonies. Colours as in Figure 1. Scalebar = ∼1 μm. (Q-U) Intercellular bridges in colonial *S. rosetta* exhibit a high diversity of morphologies, suggestive of disconnection. In addition to prior descriptions of intercellular bridges (arrowheads) and electron-dense septa (asterisks), bridges in colonial *S. rosetta* often display an asymmetrically distributed septum (Q), protracted and elongated morphology (R), disconnection from one of the contiguous cells (S) and evidence of abscission (T) and putative inheritance of the septum (U). Scalebar = 200 nm.

In animal cell types, fusion/fission dynamics have been previously associated with cellular stress [17] and substrate availability [18], but it is of most interest for choanoflagellates in the context of aerobic metabolism. For example, the fresh water choanoflagellate *Desmarella moniliformis* exhibits a shift in mitochondrial profile prior to encystment and metabolic dormancy [19] and choanoflagellates have been uncovered from hypoxic waters [20]. The role of oxygen in the origin and evolution of animals has long been discussed [21] and is currently met with controversy [22,23]. Coupled to a previous report of positive aerotaxis in *S. rosetta* rosette colonies [24], this finding places even more emphasis on understanding variation in aerobic metabolism between single and colonial choanoflagellates.

Finally, we found that colonial cells are characterised by a more amoeboid morphology than single cells (Figure 3A). Colonial cells exhibited a higher relative proportion of endocytotic vacuoles by volume (single: 0.07 ± 0.07 vs colonial: 0.32 ± 0.12) - a phenomenon coupled to a higher overall number of endocytotic vacuoles (single: 1 ± 1 vs colonial: 5 ± 2) and pseudopodial projections per cell (single: 1 ± 1 vs colonial 8 ± 2) (Figure 2C and Tables S1 and S2). Many of the pseudopodial projections and endocytotic vacuoles bore the morphology of lamellipod ruffles and macropinosomes (Figure 3A), suggesting that colonial cells are typified by high macropinocytotic activity. Macropinocytosis – defined as the formation of phase-lucent vacuoles >0.2 μm in diameter from wave-like, plasma membrane ruffles [25] – is conserved from the Amoebozoa [26] to animal cell types [27]. It is therefore parsimonious to infer that the macropinocytotic activity of *S. rosetta* colonial cells represents a trophic adaptation, particularly considering that previous biophysical studies have reported more favourable feeding hydrodynamics in rosette colonies [28]. Even in macropinosomes with no observable cargo, dissolved proteins [29] and ATP [27] from extracellular fluid have been previously reported to be metabolically exploited by animal macropinocytotic cell types. This non-selectivity, coupled to the large volume of engulfed fluid, makes macropinocytosis an efficient cellular process to sample and process the extracellular milieu.

Our comparison between single and colonial cells provides new insights into ultrastructural commonalities and differences associated with the conversion from solitary to colonial cells and shows that colonial cells might represent a distinct and differentiated cell type.

### Reconstruction of multiple rosettes reveals colony-wide cell arrangement, different cell shapes and complete cell-cell contact network

While high magnification 3D ssTEM enabled the high-resolution reconstruction of individual colonial cells, their context and interactions with neighbouring cells were lost. To address this, we reconstructed the subcellular structures of a seven-cell rosette colony (complete rosette, RC1) from 80 nm sections taken at lower magnification (Figure 3A-D, Video S7), as well as the gross morphology of four larger rosettes (RC2-5) from 150 nm sections to provide a more representative survey (Figure 3E-P).

We found that individual cells in rosette colonies vary widely in volume (Figure 3M, N), although no pattern was detected in the volumetric cellular arrangement along the rosette z-axis (Figure 3M). In addition, mean cell size was comparable among different rosettes, including those that contained different numbers of cells (Figure S4B). However, we did find a positive correlation between cell number and the number of intercellular bridges per cell across rosette colonies (Figure S4B).

Importantly, we uncovered the presence of unusually shaped cells in two of the five *S. rosetta* rosette colonies (Carrot-shaped cell 5 in RC3 and chili-shaped cell 5 in RC4, both labelled orange with an asterisk) (Figure 3M). These unusual cells were both found at the same location along the rosette z-axis, exhibited an elongated morphology distinct from other colonial cells (Figure 3O, P and Videos S8 and S9), and were small in volume. Cells 5 from RC3 and RC4 were 9.87 and 13.35 μm^3^ respectively (Figure 3N) - the mean volume of the cells in RC3 and RC4 was 27.38 and 27.25 μm^3^ respectively (Figure 3N). While each of these unusual cells possessed a flagellum, a collar, connections to neighbouring cells via intercellular bridges and had a similar proportion of cell volume dedicated to most of their major organelles as observed in other colonial cells, these cells devoted a larger volumetric percentage of the cell body to the nucleus (29.8% and 30.78% respectively versus the mean colonial proportion of 13.76 ± 0.49%). These data hint that cell differentiation within colonies may be more complex than previously realized.

Our 3D ssTEM reconstructions of rosette colonies also revealed the distribution of intercellular bridges, and the connections formed between individual cells (Figure 3M). We found intercellular bridges in all analysed rosette colonies (RC1-5), totalling 36 bridges. There was no detectable pattern regarding bridge networking across rosette colonies. Bridges were distributed from the cell equator to either of the poles along the cellular z-axis and the average bridge was 0.75 ± 0.38 μm in length (Figure S4D). Prior studies [3,4] of *S. rosetta* bridges suggested that bridges are typically short (0.15 μm), connecting two adjacent cells and containing parallel plates of electron-dense material. In contrast, the bridges detected in this study exhibited striking morphological diversity (Figure 3M, Q-U), with lengths ranging from 0.21 – 1.72 μm. The majority of bridges consisted of a protracted cytoplasmic connection between two cells, and in many cases, the septum was localized asymmetrically along the bridge (Figure S4C). Most surprisingly, some bridges were not connected to any neighbouring cells at all, but rather the septum was situated on the end of a thin, elongated cellular protrusion (Figure 3S). In addition, we observed asymmetric bridge width and degraded electron dense structures proximal to bridge remnants being incorporated into the cell body of a contiguous cell (Figure 3T, U). These data suggest that intercellular bridges could be disconnected from neighbouring cells and that the electron-dense septum may be inherited.

The asymmetric and disconnected morphology of intercellular bridges provides important clues to choanoflagellates colony formation and potentially the evolution of animal multicellularity. Bridges, displaying electron-dense septa reminiscent of those found in *S. rosetta*, have been previously identified in other colony-forming choanoflagellate species [30,31] and it has been hypothesized that these structures represent stable channels for intercellular communication [4]. Our data suggest that bridges can be disconnected, and that the electron-dense septum may be asymmetrically inherited. In this way, choanoflagellate bridges may resemble the mitotic midbody in animal cells [32]. It may still be that *S. rosetta* bridges play a role in cell-cell communication, albeit transiently. However, the exit of colonial cells from the rosette (as previously reported [3]) must involve bridge disconnection, and a proper understanding of the fate of the septum could augment our understanding of choanoflagellate cell differentiation and destiny in colony development.

### Three-dimensional cellular architecture of sponge choanocytes

To place our choanoflagellate reconstructions into the context of collar cells from an early-branching animal, we reconstructed a section of a sponge choanocyte chamber (Figure 4A) from the homoscleromorph sponge *Oscarella carmela* [33]. Both choanoflagellates and sponge collar cells influence local hydrodynamics by beating their single flagellum to draw in bacteria that are captured by the apical collar complex [34], however sponge choanocytes are part of an obligately multicellular organism. Our 3D ssTEM reconstructions allowed for the reconstruction of five choanocytes and for the volumetric and numerical comparison of choanocyte and choanoflagellate subcellular structures (Figure 4B-E, Figures S5, S6 and Video S10). We detected little ultrastructural variability within the five choanocytes (Figure S5, Table S3 and S4). All five cells exhibited a prominent basal nucleus, small and unreticulated mitochondria, food vacuoles scattered around the entire cell, and an apical Golgi apparatus (Figure 4B-D and Figure S5, S6) - consistent with the coarse choanocyte cellular architecture reported in previous studies [32,34,36] (reviewed in [1,37]).

**Figure 4.**
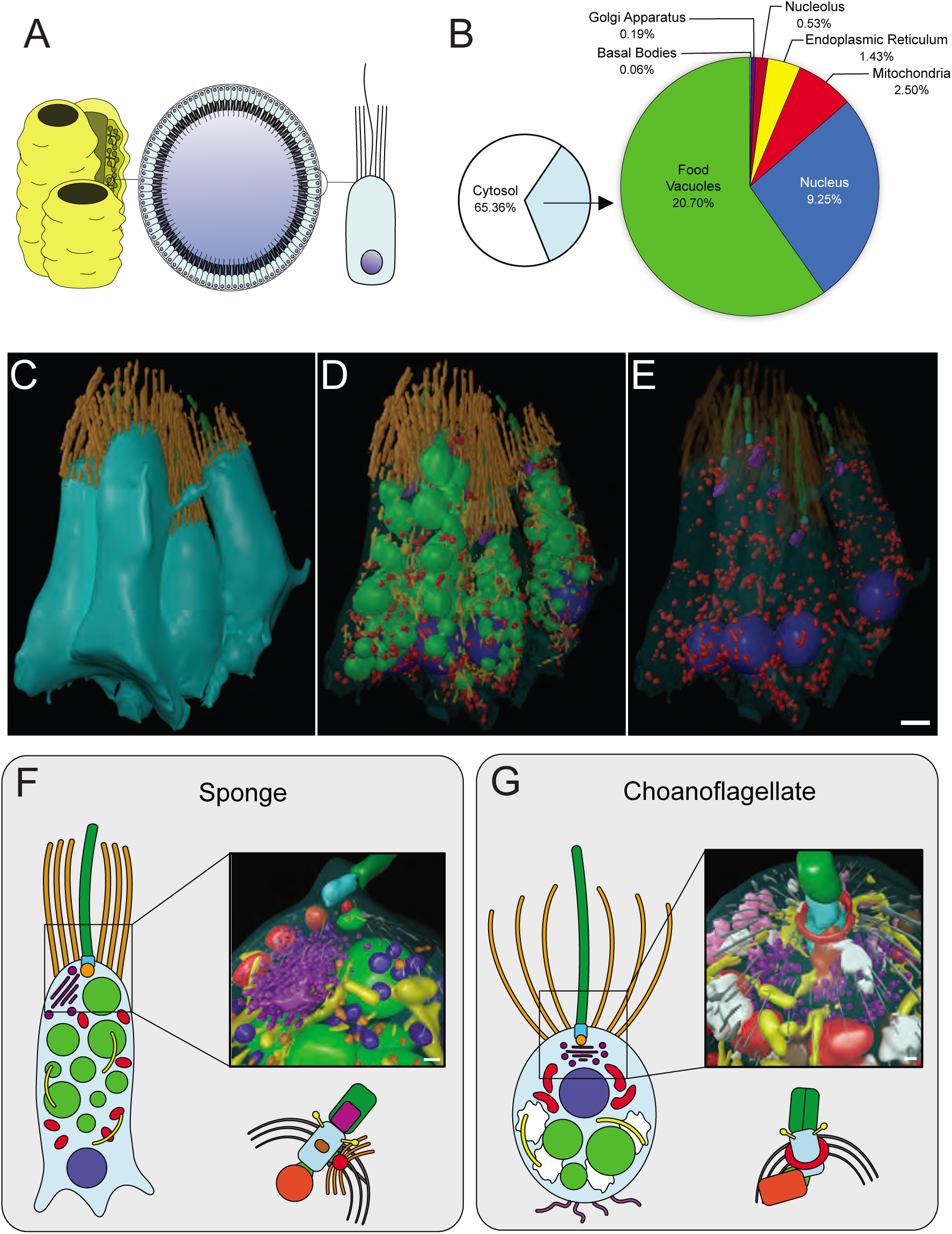
Three-dimensional cellular architecture of sponge choanocytes. (A) Choanocytes line interconnected chambers in members of the Porifera and serve as feeding cells. (B) Mean volumetric breakdown of five sponge choanocytes. Colours are as in Figure 1. (C-E) 3D ssTEM of a section of choanocyte chamber containing five complete cells (B). The plasma membrane was rendered transparent (D) and food vacuoles and ER were removed to allow better visualisation of subcellular structures (E). Colours are as in Figure 1. Scalebar = ∼1 μm. (F-G) Reconstruction and comparison of the sponge choanocyte (F) and choanoflagellate (G) apical poles shows distinct differences between the two cell types. Shown in the choanocyte reconstruction are the basal foot (red, associated with basal body), food vacuole (light green), endoplasmic reticulum (yellow) flagellar basal body (light blue), flagellum (dark green), Golgi apparatus and Golgi-associated vesicles (purple), microtubules (grey), mitochondria (red)m non-flagellar basal body (dark orange), Type 1 vesicles (light orange) and Type 2 vesicles. Shown in the choanoflagellate reconstruction are the apical vesicles (pink), food vacuole (light green), endoplasmic reticulum (yellow) flagellar basal body (light blue), flagellum (dark green), Golgi apparatus and Golgi-associated vesicles (purple), glycogen (white), large vesicles (brown), microtubules (grey), microtubular ring (red) and non-flagellar basal body (dark orange). Scalebars = 200 nm. Diagrams of the choanocyte fine kinetid (F) and choanoflagellate fine kinetid (G) structure highlight the distinct differences.

Furthermore, our data showed many ultrastructural commonalities between choanocytes and choanoflagellates. For example, the number of microvilli that surround the apical flagellum in single and colonial choanoflagellates is comparable to the number of microvilli in sponge choanocytes (single: 32 ± 2 vs colonial: 35.3 ± 4.9 vs choanocytes: 30.6 ± 4.1) (Figure S6A). We also found that the number of food vacuoles and the number and volumetric proportion of the Golgi apparatus are similar in all three cell types (Figure S6A). Although, choanocytes did not appear to exhibit the same macropinocytotic activity as colonial choanoflagellates throughout the cell (some micropinocytotic inclusions are present towards the cell apex (Figure S6 D-E)), basal sections of choanocytes were heavily amoeboid (Figure S6 B-C). These amoeboid protrusions may not only be for mechanical anchorage into the mesohyl, but may play a role in phagocytosis as we observed bacteria in the mesohyl to be engulfed by basal pseudopodia (Figure S6 F-G). Thus, both choanocytes and colonial choanoflagellates are typified by high amoeboid cell activity.

Not unexpectedly, we also observed some ultrastructural differences between choanocytes and choanoflagellates. In contrast with cells from choanoflagellate rosettes, sponge choanocytes lack filopodia and intercellular bridges. Choanocytes also do not possess glycogen reserves and devote significantly less of their cell volume (9.25 ± 0.39%) than choanoflagellates (single: 12.92 ± 0.58% and colonial: 11.56 ± 0.27%) to the nucleus, and less to mitochondria (2.5% ± 0.3% versus single: 5.08 ± 1.14% and colonial: 6.63 ± 0.42%) (Figure S6A). However, choanocytes devote significantly more of their volume to food vacuoles (20.7 ± 1.01%) than choanoflagellates (single: 9.22 ± 2.75% and colonial: 6.85 ± 0.87%) (Figure 4E). High-resolution reconstructions of the choanocyte and choanoflagellate apical pole (Figure 4 F-G and Videos S11, S12) showed differences in terms of vesicle type and localisation, Golgi positioning and collar arrangement (conical in choanoflagellates while cylindrical in choanocytes, as previously noted [34]). The flagellar basal body has previously been meticulously characterised in both choanocytes and choanoflagellates and some differences have been reported between the two by other authors [38–43] These findings are reiterated by our reconstructions and observations (Figure 4F, G).

### Concluding Remarks

The comparative 3D reconstruction of collar cells from two different phyla, choanoflagellates and sponges, allowed for an unbiased view of their cellular architecture and for the reconstruction of key properties of the enigmatic ancestral collar cell. Our data reveal distinct ultrastructural features in single and colonial choanoflagellates and demonstrate that cells within rosette colonies vary significantly in their cell size and shape. The newly identified ‘carrot’ and ‘chili cells’ reveal that cells within choanoflagellate colonies do not simply consist of an assemblage of equivalent single cells but some may represent a distinctly differentiated cell type displaying ultrastructural modifications. Likewise, our data suggest that sponge choanocytes are not simply an incremental variation of the choanoflagellate cell, but are specialised feeding cells as indicated by their high volumetric proportion of food vacuoles. Together, our data show a remarkable variety of collar cell architecture and suggest cell type differentiation was present in the stem lineage leading to animals.

## ACKNOWLEDGEMENTS

The authors would like to thank Charlotte Walker and Glen Wheeler for their technical support with fluorescent microscopy, and generously providing the vital stains. We thank Scott Nichols for providing *Oscarella carmela* samples and Pete Bond from the Plymouth Electron Microscopy Lab for his valuable assistance during the imaging of the three high-resolution choanoflagellate colonial cells. Finally, we are grateful to Manuela Truebano Garcia for her continued advice and feedback throughout this project. This work was supported by the Anne Warner endowed Fellowship through the Marine Biological Association of the UK, the Royal Society University Research Fellowship and Sars core budget.

## AUTHOR CONTRIBUTIONS

DL, KM, PB designed the study; DL, BL, KM, PB performed experiments; DL, KM, PB analysed data; DL, NK, PB wrote the paper and all authors reviewed, commented on, and edited the manuscript.

## DECLARATION OF INTRESTS

The authors declare no competing financial interest

## FIGURE LEGENDS

**Figure S1.**
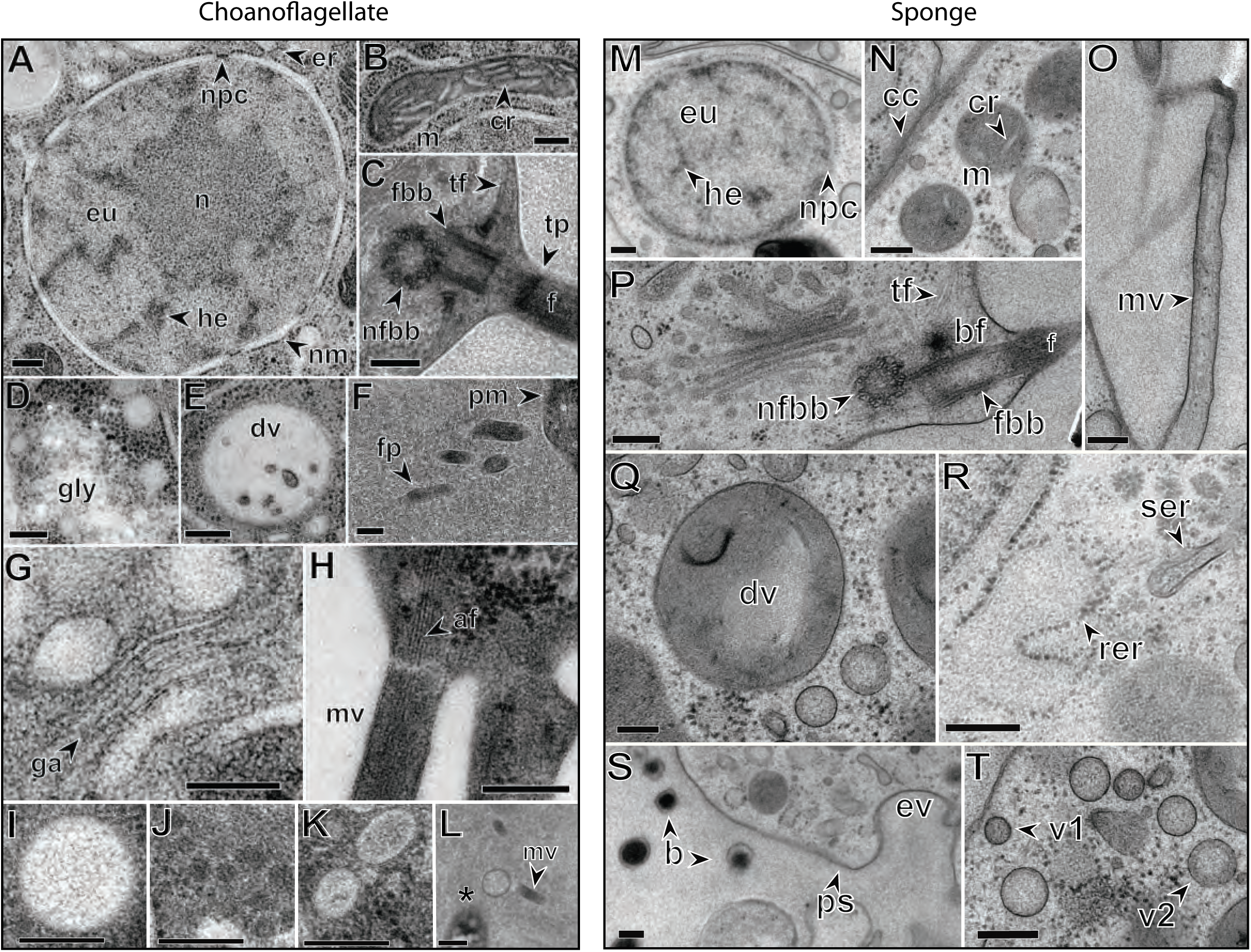
High magnification TEM panel of the *S. rosetta* (A-L) and *O. carmela* (M-T) subcellular components discussed herein. (A) *S. rosetta* nucleus showing endoplasmic reticulum (er), euchromatin (eu), heterochromatin (he), nuclear membrane (nm), nuclear pore complex (npc) and nucleolus (n). (B) Mitochondrion (m) showing flattened, non-discoidal cristae (cr). (C) Apical pole showing flagellum (f), flagellar basal body (fbb), non-flagellar basal body (nfbb), tubulin filaments (tf) and transversal plate (tp). (D) Area of high glycogen storage (gly). (E) Food vacuole (dv). (F) Posterior filopodia (fp) projecting from the basal plasma membrane (pm). (G) Golgi apparatus (ga). (H) microvillus (mv) from the apical collar displaying actin filaments (af). (I) Large, extremely electron-lucent vesicles. (J) Golgi-associated, electron dense vesicles. (K) Apical, electron-lucent vesicles. (L) Extracellular vesicles were observed in two of the single cells and appeared to bud from the microvillar membrane. (M) *O. carmela* nucleus showing euchromatin (eu), heterochromatin (he) and nuclear pore complex (npc). (N) Mitochondria (m) displaying cristae (cr). Also visible are cell-cell contacts between two adjacent choanocytes (cc). (O) Collar microvillus (mv). (P) Apical pole and Golgi apparatus showing flagellum (f), flagellar basal body (fbb), non-flagellar basal body (nfbb), tubulin filaments (tf) and basal foot (bf). (Q) Food vacuole (dv). (R) Rough (rer) and smooth (ser) endoplasmic reticulum. (S) Basal pole of *O. carmela* shows bacteria located in the mesohyl (b), basal pseudopodia (ps) and endocytotic invagination (ev). (T) Vesicles type 1 (V1) and type 2 (V2) are located throughout the choanocyte cytoplasm. Scale bars = 200 nm.

**Figure S2.**
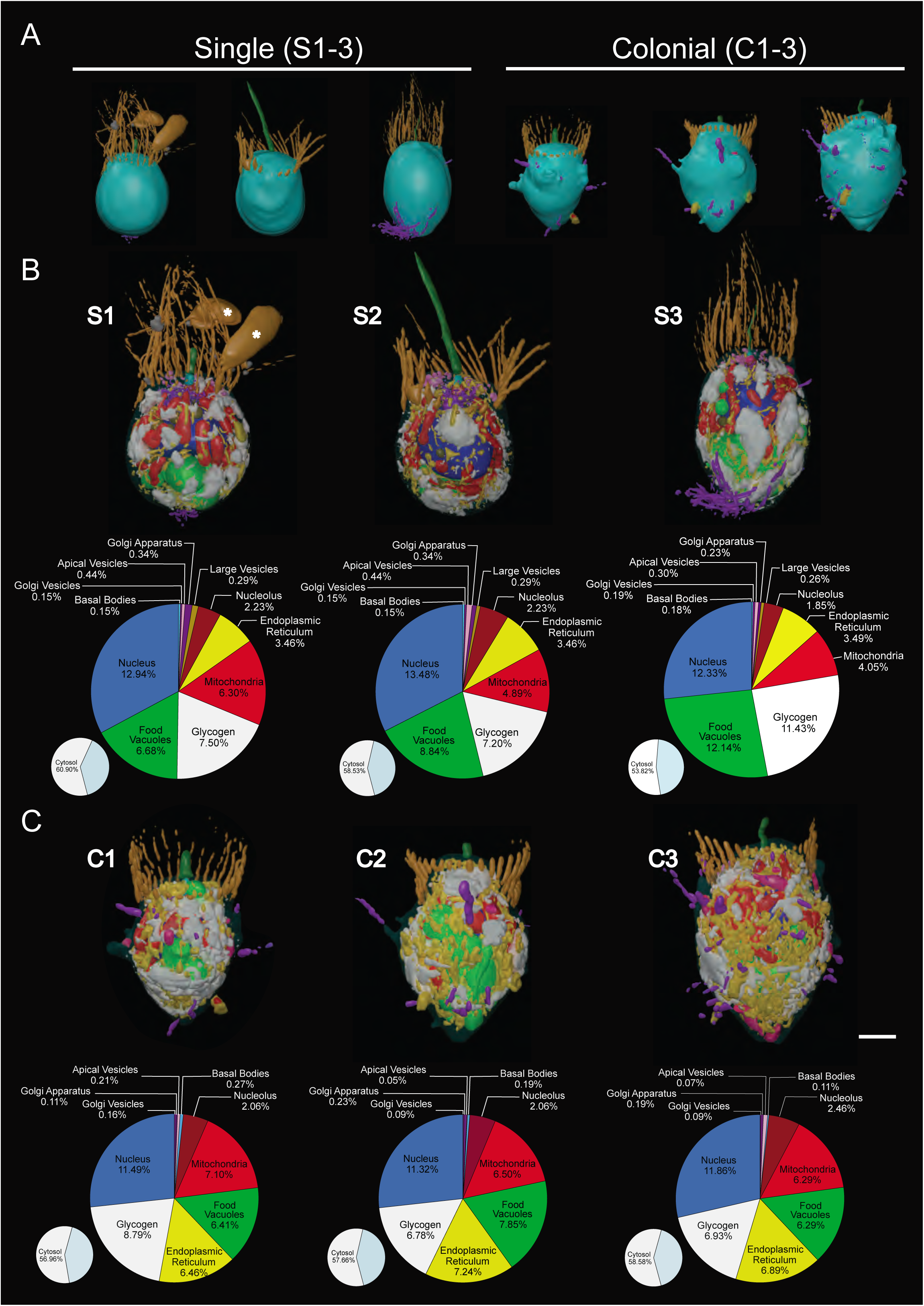
3D ssTEM reconstructions of high resolution single and colonial *S. rosetta* cells. (A) Gross external morphologies of reconstructions of both single (S1-3) and colonial (C1-3) *S. rosetta* cells. (B-C) Structomic reconstructions of single (B) and colonial (C) *S. rosetta* cells, with the plasma membrane removed to reveal subcellular ultrastructure. Colours are as in Figure 1. Asterisks indicate engulfed prey bacteria. Cells are labelled with their corresponding cell ID number and volumetric breakdown for each cell is shown below reconstructions. Scalebar = ∼1 μm.

**Figure S3.**
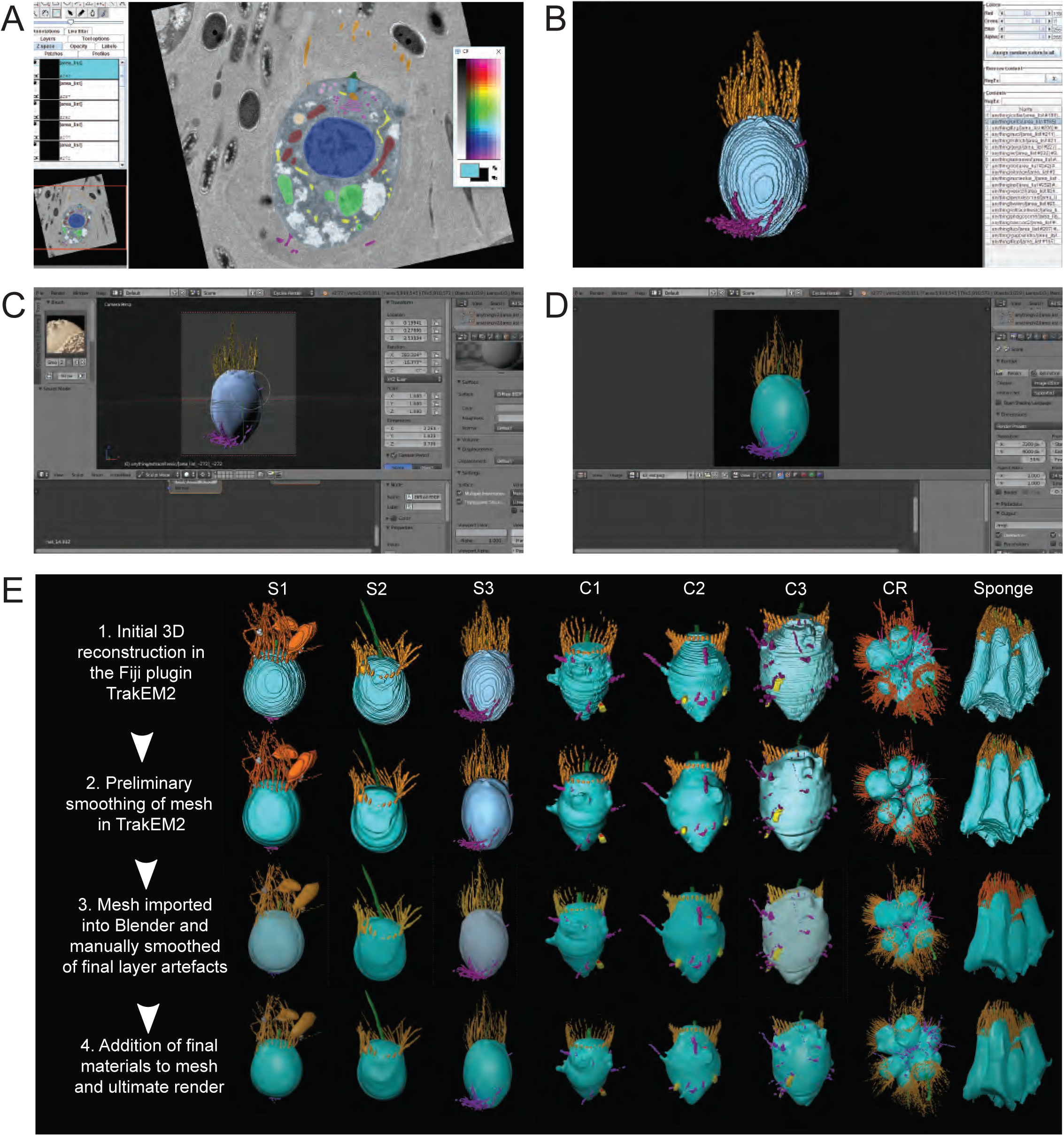
Methodological overview of 3D ssTEM reconstruction of *S. rosetta* and *O. carmela* cells. (A) ssTEM stacks are imported into the Fiji plugin TrakEM2, aligned, and scaled. Subcellular structures are then manually segmented. (B) 3D ssTEM reconstructions are conducted in TrakEM2 by merging traced structures along the z-axis, initially smoothed and imported into Blender (C). In Blender, final reconstruction artefacts are smoothed using the F Smooth Sculpt Tool and final materials are added for the ultimate render (D). (E) The aforementioned methodology applied to single cells (S1-3), colonial cells (C1-3), a complete rosette colony (CR) and a section of an *O. carmela* choanocyte chamber

**Figure S4.**
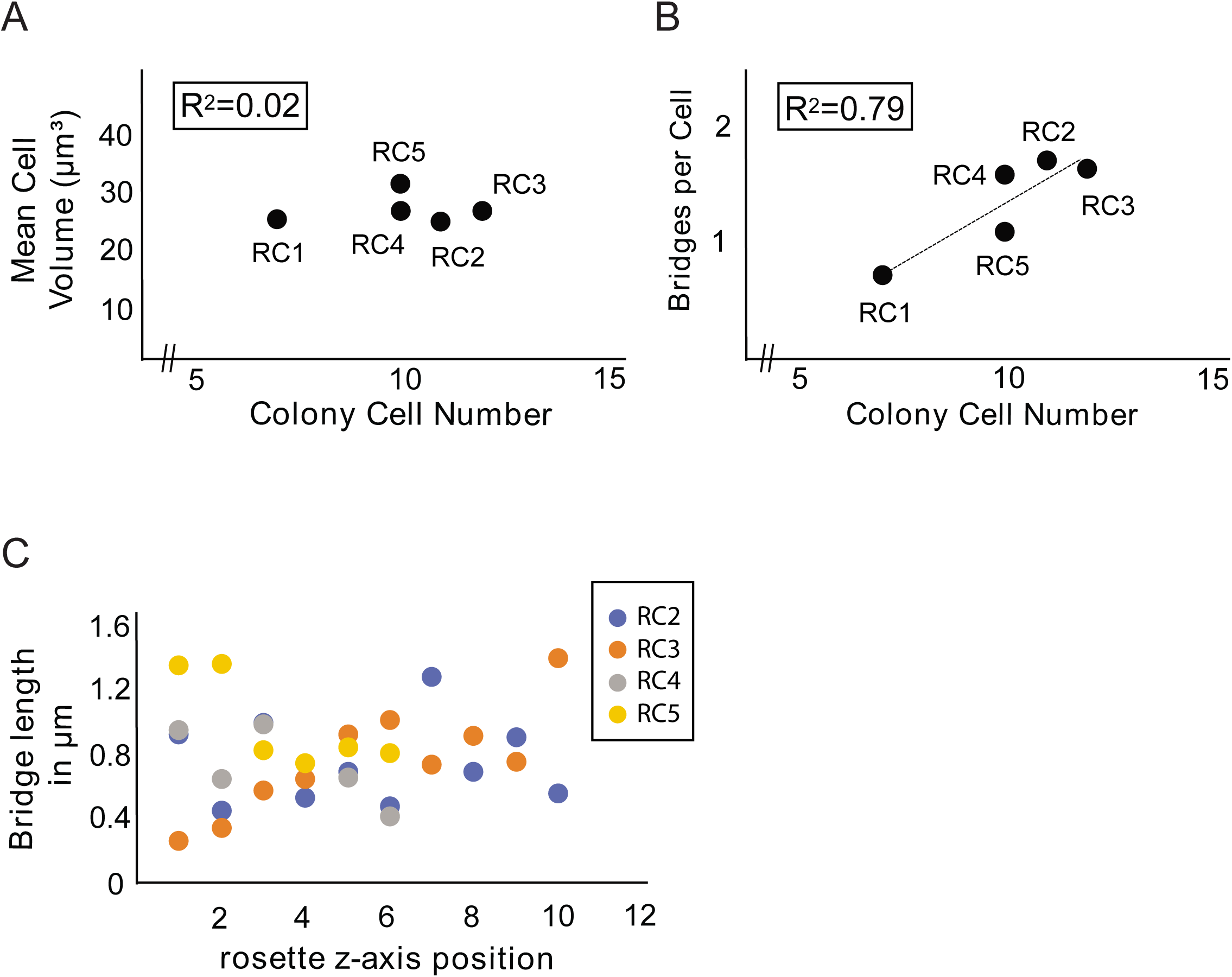
Mean cell volume per colony cell number, intercellular bridges per colony cell number and bridge length. (A) No correlation was found between cell volume and colony cell number. (B) A positive correlation was found between bridges per cell and colony cell number (*p*<0.05). (C) No apparent pattern was observed between the length of an intercellular bridge and its position along the colony z-axis.

**Figure S5.**
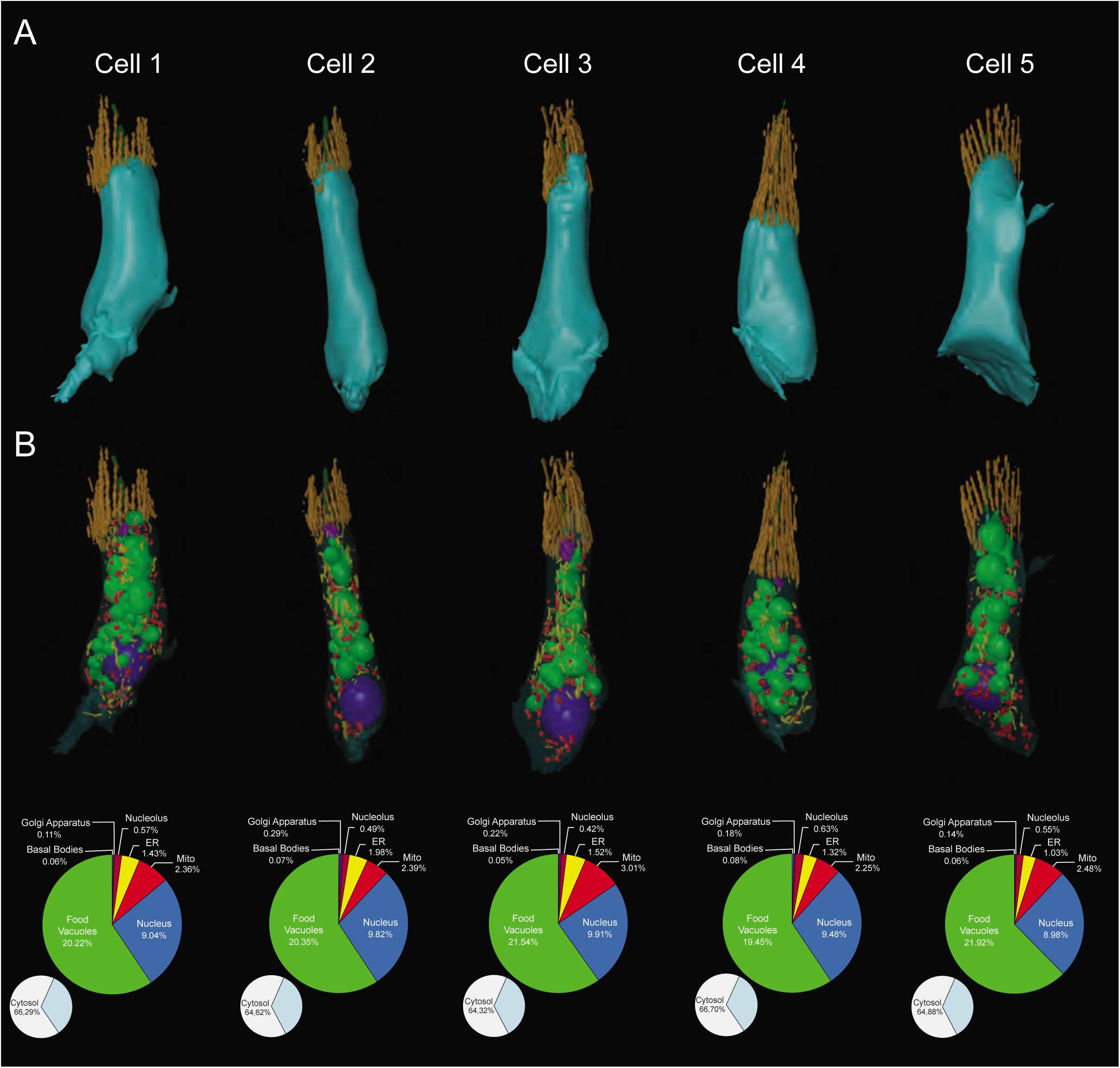
3D reconstructions and volumetric breakdown of five sponge choanocytes. (A-B) 3D ssTEM reconstructions of five *O. carmela* choanocytes and their volumetric breakdown is shown below. Scalebar = ∼1 μm.

**Figure S6.**
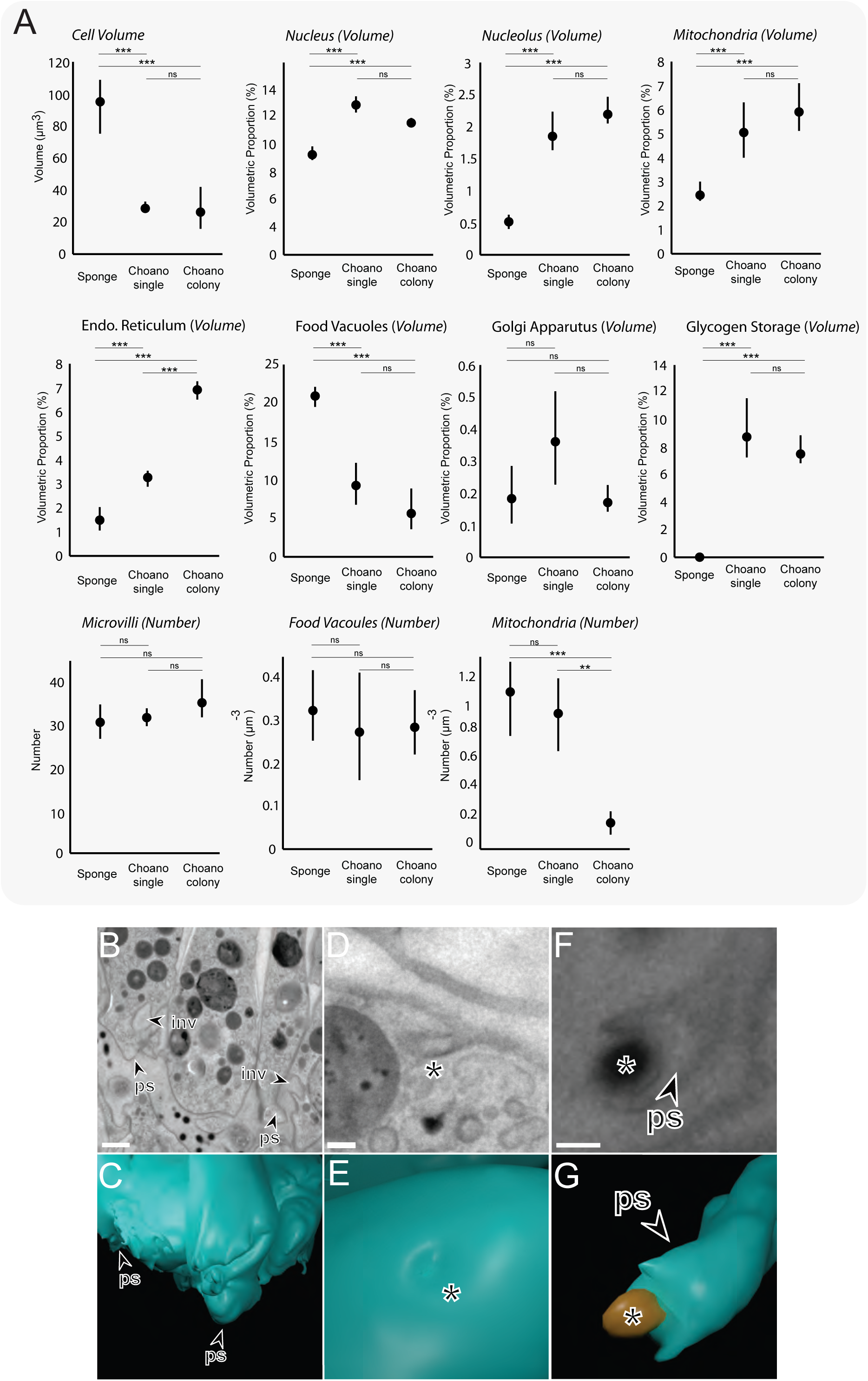
Volumetric and numerical comparison of choanocyte and choanoflagellate major subcellular structures. (A) Choanocytes from *O. carmela* are significantly larger by volume (μm^3^) than the single and colonial choanoflagellate *S. rosetta* cells. Volumetric (%) (±SEM) (nucleus, nucleolus, mitochondria, endoplasmatic reticulum, food vacuoles and glycogen storage) and numerical (μm^-3^) (±SEM) (mitochondria) differences were found between sponge choanocytes (*n* = 5) and single (n = 3) and colonial (n = 3) choanoflagellates. * *p* < 0.05, ** *p* < 0.01, *** *p* < 0.001. (B-G) TEM and 3D ssTEM reconstructions of amoeboid cell behaviour in sponge choanocytes. Shown are the highly invaginated (inv) and pseudopodiated (ps) basal pole of the choanocyte (B&C), macropinocytotic activity (*) at the apical pole (D&E) and a mesohyl-associated bacterium being engulfed by a pseudopodium (ps) at the basal pole (F&G).

**Table S1.** Volumetric measurements of *S. rosetta* cells and components.

|  | Single cells |  |  |  | Colonial cells |  |  |  |
| --- | --- | --- | --- | --- | --- | --- | --- | --- |
| **Organelle** | **S1** | **S2** | **S3** | **Mean +/- SD** | **C1** | **C2** | **C3** | **Mean +/- SD** |
| Cell Body | 26.81<br>(100) | 32.34<br>(100) | 26.68<br>(100) | 28.61 ± 3.23<br>(100 ± 0) | 18.89<br>(100) | 21.54<br>(100) | 41.82<br>(100) | 27.41 ± 12.54<br>(100 ± 0) |
| Nucleus | 3.92<br>(12.94) | 5.08<br>(13.48) | 3.73<br>(12.33) | 4.24 ± 0.73<br>(12.92 ± 0.58) | 2.56<br>(11.49) | 2.89<br>(11.33) | 5.99<br>(11.86) | 3.81 ± 1.89<br>(11.56 ± 0.27) |
| Nucleolus | 0.45<br>(1.68) | 0.72<br>(2.23) | 0.44<br>(1.65) | 0.54 ± 0.16<br>(1.85 ± 0.33) | 0.39<br>(2.06) | 0.45<br>(2.09) | 1.03<br>(2.46) | 0.62 ± 0.35<br>(2.2 ± 0.22) |
| Mitochondria | 1.69<br>(6.30) | 1.58<br>(4.89) | 1.08<br>(4.05) | 1.45 ± 0.33<br>(5.08 ± 1.14) | 1.34<br>(7.10) | 1.4<br>(6.50) | 2.63<br>(6.29) | 1.79 ± 0.73<br>(6.63 ± 0.42) |
| Endoplasmic Reticulum | 0.77<br>(2.87) | 1.12<br>(3.46) | 0.93<br>(3.49) | 0.94 ± 0.18<br>(3.27 ± 0.35) | 1.22<br>(6.46) | 1.56<br>(7.24) | 2.88<br>(6.89) | 1.89 ± 0.88<br>(6.86 ± 0.39) |
| Food Vacuoles | 1.79<br>(6.68) | 2.86<br>(8.84) | 3.24<br>(12.14) | 2.63 ± 0.75<br>(9.22 ± 2.75) | 1.21<br>(6.41) | 1.69<br>(7.85) | 2.63<br>(6.29) | 1.84 ± 0.72<br>(6.85 ± 0.87) |
| Glycogen Storage | 2.01<br>(7.50) | 2.33<br>(7.20) | 3.05<br>(11.43) | 2.46 ± 0.53<br>(8.71 ± 2.36) | 1.66<br>(8.79) | 1.46<br>(6.78) | 2.9<br>(6.93) | 2.01 ± 0.78<br>(7.50 ± 1.12) |
| Flagellar Basal Body | 0.02<br>(0.07) | 0.03<br>(0.09) | 0.01<br>(0.10) | 0.02 ± 0.01<br>(0.09 ± 0.02) | 0.03<br>(0.16) | 0.03<br>(0.14) | 0.02<br>(0.05) | 0.03 ± 0.01<br>(0.12 ± 0.06) |
| Non-Flagellar Basal Body | 0.02<br>(0.07) | 0.02<br>(0.06) | 0.02<br>(0.08) | 0.02 ± 0<br>(0.07 ± 0.01) | 0.02<br>(0.11) | 0.01<br>(0.05) | 0.02<br>(0.05) | 0.02 ± 0.01<br>(0.07 ± 0.03) |
| Golgi Apparatus | 0.14<br>(0.52) | 0.11<br>(0.34) | 0.06<br>(0.23) | 0.10 ± 0.04<br>(0.36 ± 0.15) | 0.02<br>(0.11) | 0.05<br>(0.23) | 0.08<br>(0.19) | 0.05 ± 0.03<br>(0.18 ± 0.06) |
| Golgi Associated Vesicles | 0.03<br>(0.12) | 0.05<br>(0.15) | 0.05<br>(0.19) | 0.04 ± 0.01<br>(0.15 ± 0.04) | 0.03<br>(0.16) | 0.02<br>(0.09) | 0.03<br>(0.07) | 0.03 ± 0.01<br>(0.11 ± 0.05) |
| Apical Vesicles | 0.06<br>(0.22) | 0.14<br>(0.44) | 0.08<br>(0.30) | 0.09 ± 0.04<br>(0.32 ± 0.11) | 0.04<br>(0.21) | 0.01<br>(0.05) | 0.14<br>(0.33) | 0.06 ± 0.07<br>(0.20 ± 0.14) |
| Large Vesicles | 0.03<br>(0.45) | 0.09<br>(0.29) | 0.07<br>(0.26) | 0.06 ± 0.03<br>(0.33 ± 0.10) | 0<br>(0) | 0<br>(0) | 0<br>(0) | 0<br>(0) |
| Extracellular Vesicles | 0.06<br>(0.22) | 0<br>(0) | 0.01<br>(0.04) | 0.02 ± 0.03<br>(0.09 ± 0.12) | 0<br>(0) | 0<br>(0) | 0<br>(0) | 0<br>(0) |
| Endocytotic Vacuoles | 0.04<br>(0.15) | 0<br>(0) | 0.02<br>(0.07) | 0.02 ± 0.02<br>(0.07 ± 0.07) | 0.06<br>(0.32) | 0.04<br>(0.19) | 0.18<br>(0.43) | 0.09 ± 0.08<br>(0.32 ± 0.12) |
| Filopodia | 0.07<br>(0.26) | 0<br>(0) | 0.28<br>(1.05) | 0.12 ± 0.15<br>(0.44 ± 0.55) | 0.07<br>(0.37) | 0.08<br>(0.37) | 0.18<br>(0.43) | 0.11 ± 0.06<br>(0.39 ± 0.03) |
| Cytosol | 16.33<br>(60.9) | 18.93<br>(58.53) | 14.36<br>(53.82) | 16.54 ± 2.29<br>(57.75 ± 3.6) | 10.76<br>(56.96) | 12.42<br>(57.66) | 24.50<br>(58.58) | 15.89 ± 7.49<br>(57.97 ± 0.81) |

Volumes were measured in $\mu\text{m}^3$ .
Values between parentheses are percentages of cell volume.

**Table S2.**
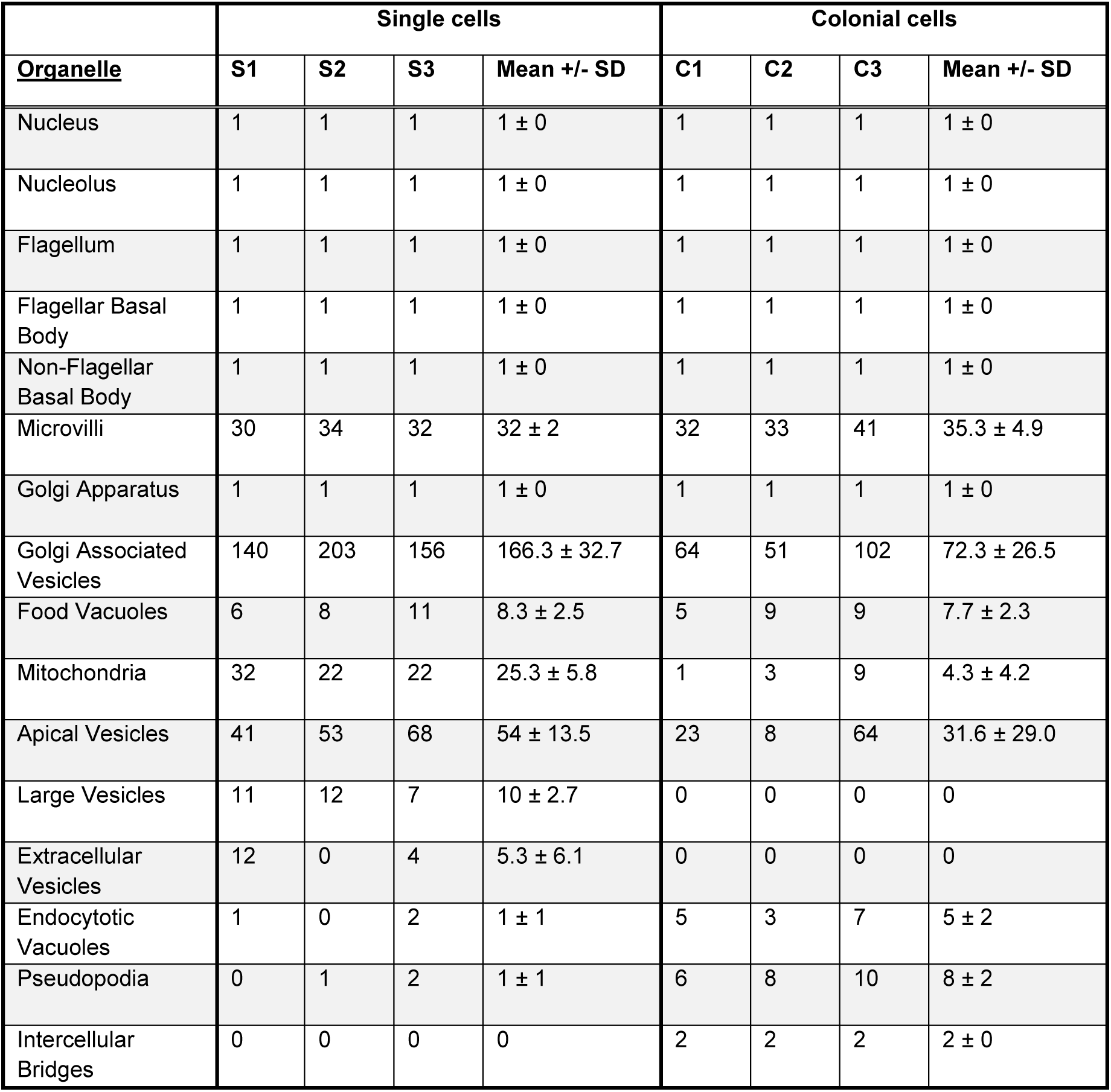
Numbers of various organelles and components in *S. rosetta* cells.

**Table S3.**
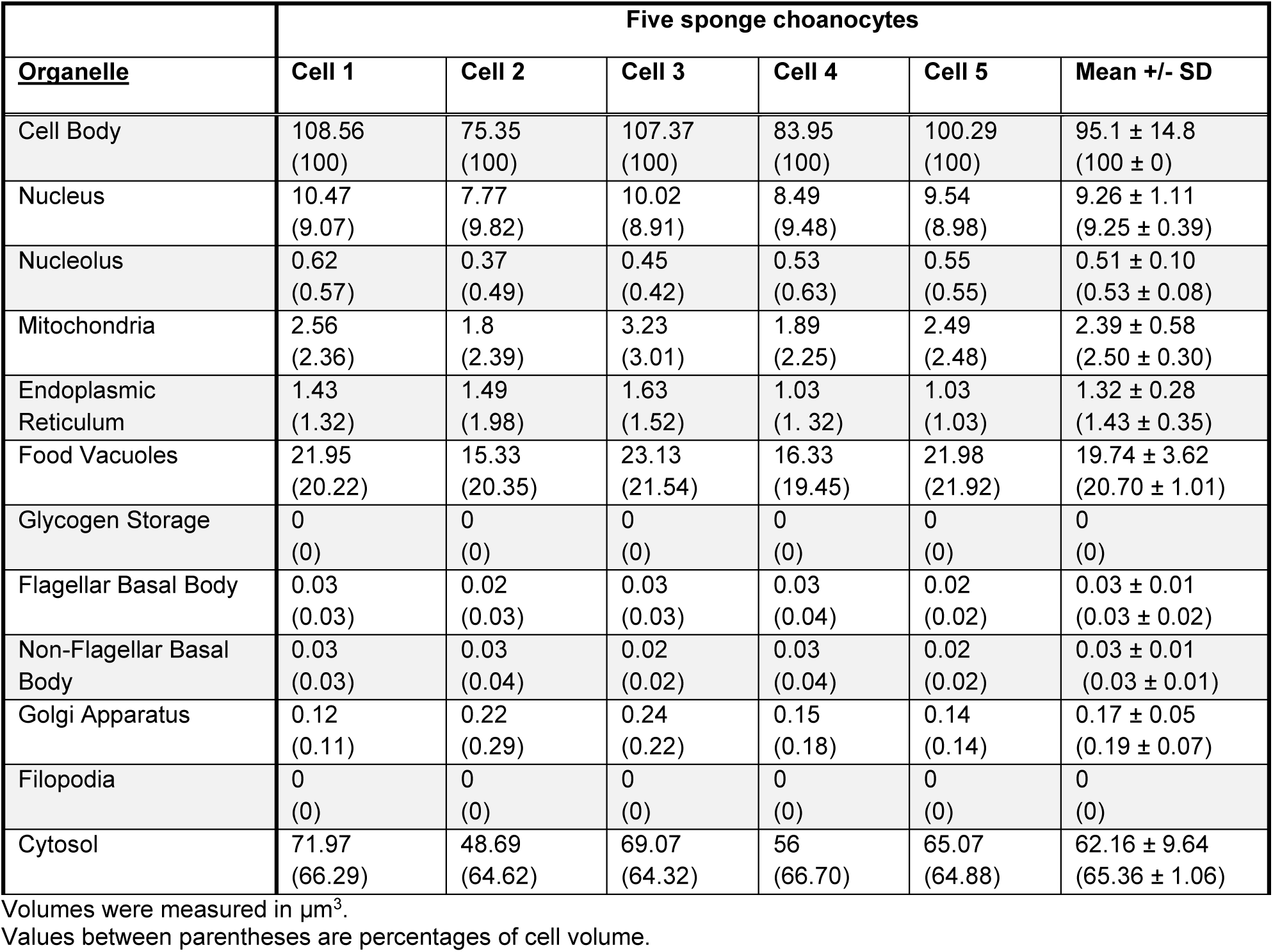
Volumetric measurements of *O. carmela* choanocytes and components.

|  | Five sponge choanocytes |  |  |  |  |  |
| --- | --- | --- | --- | --- | --- | --- |
| <u>Organelle</u> | Cell 1 | Cell 2 | Cell 3 | Cell 4 | Cell 5 | Mean +/- SD |
| Cell Body | 108.56<br>(100) | 75.35<br>(100) | 107.37<br>(100) | 83.95<br>(100) | 100.29<br>(100) | 95.1 ± 14.8<br>(100 ± 0) |
| Nucleus | 10.47<br>(9.07) | 7.77<br>(9.82) | 10.02<br>(8.91) | 8.49<br>(9.48) | 9.54<br>(8.98) | 9.26 ± 1.11<br>(9.25 ± 0.39) |
| Nucleolus | 0.62<br>(0.57) | 0.37<br>(0.49) | 0.45<br>(0.42) | 0.53<br>(0.63) | 0.55<br>(0.55) | 0.51 ± 0.10<br>(0.53 ± 0.08) |
| Mitochondria | 2.56<br>(2.36) | 1.8<br>(2.39) | 3.23<br>(3.01) | 1.89<br>(2.25) | 2.49<br>(2.48) | 2.39 ± 0.58<br>(2.50 ± 0.30) |
| Endoplasmic Reticulum | 1.43<br>(1.32) | 1.49<br>(1.98) | 1.63<br>(1.52) | 1.03<br>(1.32) | 1.03<br>(1.03) | 1.32 ± 0.28<br>(1.43 ± 0.35) |
| Food Vacuoles | 21.95<br>(20.22) | 15.33<br>(20.35) | 23.13<br>(21.54) | 16.33<br>(19.45) | 21.98<br>(21.92) | 19.74 ± 3.62<br>(20.70 ± 1.01) |
| Glycogen Storage | 0<br>(0) | 0<br>(0) | 0<br>(0) | 0<br>(0) | 0<br>(0) | 0<br>(0) |
| Flagellar Basal Body | 0.03<br>(0.03) | 0.02<br>(0.03) | 0.03<br>(0.03) | 0.03<br>(0.04) | 0.02<br>(0.02) | 0.03 ± 0.01<br>(0.03 ± 0.02) |
| Non-Flagellar Basal Body | 0.03<br>(0.03) | 0.03<br>(0.04) | 0.02<br>(0.02) | 0.03<br>(0.04) | 0.02<br>(0.02) | 0.03 ± 0.01<br>(0.03 ± 0.01) |
| Golgi Apparatus | 0.12<br>(0.11) | 0.22<br>(0.29) | 0.24<br>(0.22) | 0.15<br>(0.18) | 0.14<br>(0.14) | 0.17 ± 0.05<br>(0.19 ± 0.07) |
| Filopodia | 0<br>(0) | 0<br>(0) | 0<br>(0) | 0<br>(0) | 0<br>(0) | 0<br>(0) |
| Cytosol | 71.97<br>(66.29) | 48.69<br>(64.62) | 69.07<br>(64.32) | 56<br>(66.70) | 65.07<br>(64.88) | 62.16 ± 9.64<br>(65.36 ± 1.06) |

Volumes were measured in $\mu\text{m}^3$ .
Values between parentheses are percentages of cell volume.

**Table S4.** Numbers of various organelles and components in *O. carmela* choanocytes.

|  | Five sponge choanocytes |  |  |  |  |  |
| --- | --- | --- | --- | --- | --- | --- |
| <u>Organelle</u> | Cell 1 | Cell 2 | Cell 3 | Cell 4 | Cell 5 | Mean +/- SD |
| Nucleus | 1 | 1 | 1 | 1 | 1 | 1 ± 0 |
| Nucleolus | 1 | 1 | 1 | 1 | 1 | 1 ± 0 |
| Flagellum | 1 | 1 | 1 | 1 | 1 | 1 ± 0 |
| Flagellar Basal Body | 1 | 1 | 1 | 1 | 1 | 1 ± 0 |
| Non-Flagellar Basal Body | 1 | 1 | 1 | 1 | 1 | 1 ± 0 |
| Microvilli | 35 | 27 | 35 | 27 | 29 | 30.6 ± 4.1 |
| Golgi Apparatus | 1 | 1 | 1 | 1 | 1 | 1 ± 0 |
| Food Vacuoles | 42 | 22 | 27 | 35 | 27 | 30.6 ± 7.9 |
| Mitochondria | 125 | 81 | 140 | 66 | 117 | 105.8 ± 31.1 |
| Pseudopodia | 0 | 0 | 0 | 0 | 0 | 0 |
| Intercellular Bridges | 0 | 0 | 0 | 0 | 0 | 0 |

## SUPPLEMENTARY EXPERIMENTAL PROCEDURES

### Cell Culture

Colony-free *S. rosetta* cultures (ATCC 50818) were grown with co-isolated prey bacteria in 0.22 μm-filtered choanoflagellate growth medium [45] diluted at a ratio of 1:4 with autoclaved seawater. Cultures were maintained at 18°C and split 1.5:10 once a week. Colony-enriched *S. rosetta* cultures (Px1) were likewise maintained, but monoxenically cultured with the prey bacterium *Algoriphagus machipongonensis* [46] to induce rosette formation.

### Fluorescent Labelling of Organelles

To support the annotation of organelles from ssTEM sections, the microanatomy of *S. rosetta* cells was chemically characterized by fluorescent vital staining. Cells were pelleted by gentle centrifugation (500x g for 10 min at 4°C) in a Heraeus^™^ Megafuge^™^ 40R (ThermoFisher Scientific) and resuspended in a small volume of culture medium. Concentrated cell suspension (500 μl) was applied to glass-bottom dishes coated with poly-L-lysine solution (P8920, Sigma-Aldrich) and left for 10-30 min until cells were sufficiently adhered. Px1 cultures were concentrated into 100 μl of culture medium to promote the adherence of rosette colonies.

Adhered cells were incubated in 500 μl of fluorescent vital dye diluted in 0.22 μmfiltered seawater. Cells were incubated with 4.9 μM Hoechst 33342 Dye for 30 min (to label nuclei); 1 μM LysoTracker^®^ Yellow HCK-123 for 1.5 h (to label food vacuoles); 3 μM FM^®^ 1-43 Dye for 1 min (to label the plasma membrane); and 250 nM MitoTracker^®^ Red CM-H2Xros for 30 min (to label mitochondria). All vital dyes were from ThermoFisher Scientific (H3570, L12491, T35356 and M7513 respectively). Fluorescent-DIC microscopy was conducted under a 100 x oil-immersion objective lens using a Leica DMi8 epifluorescent microscope (Leica, Germany). Vital dyes were viewed by excitation at 395 nm and emission at 435-485 nm (Hoechst 33342 Dye); 470 nm and emission at 500-550 nm (LysoTracker^®^ Yellow HCK-123 & FM^®^ 1-43 Dye); and 575 nm and 575-615 nm (MitoTracker^®^ Red CM-H2Xros). Micrographs were recorded with an ORCA-Flash4.0 digital camera (Hamamatsu Photonics, Japan). All cells were imaged live. No-dye controls using only the dye solvent dimethyl sulfoxide (DMSO) (D4540, Sigma-Aldrich) were run for each wavelength to identify and control for levels of background fluorescence. Chemical fixation during vital staining and TEM sectioning was avoided where possible in this study to reduce fixation artefacts.

To visualize cell bodies, flagella, filopodia and collars adherent cells were fixed for 5 min with 1 ml 6% acetone, for 15 min with 1 ml 4% formaldehyde. Acetone and formaldehyde were diluted in artificial seawater, pH 8.0. Cells were washed gently four times with 1 ml washing buffer (100 mM PIPES at pH 6.9, 1 mM EGTA, and 0.1 mM MgSO4) and incubated for 30 min in 1 ml blocking buffer (washing buffer with 1% BSA, 0.3 % Triton X-100). Cells were incubated with primary antibodies against tubulin (E7, 1:400; Developmental Studies Hybridoma Bank) diluted in 0.15 ml blocking buffer for 1 h, washed four times with 1 ml of blocking buffer, and incubated for 1 h in the dark with fluorescent secondary antibodies (1:100 in blocking buffer, Alexa Fluor 488 goat anti mouse). Coverslips were washed three times with washing buffer, incubated with Alexa Fluor 568 Phalloidin for 15 min and washed again three times with washing buffer. Coverslips were mounted onto slides with Fluorescent Mounting Media (4 ml; Prolong Gold Antifade with DAPI, Invitrogen). Images were taken with a 100x oil immersion objective on a Leica DMI6000 B inverted compound microscope and Leica DFC350 FX camera. Images presented as z-stack maximum intensity projections.

### Electron Microscopy

#### High Pressure Freezing

Cultured *S. rosettta* single and colonial cells were concentrated by gentle centrifugation (500x g for 10 min), resuspended in 20% BSA (Bovine Serum Albumin, Sigma) made up in artificial seawater medium and concentrated again. Most of the supernatant was removed and the concentrated cells transferred to high pressure freezing planchettes varying in depth between 50 and 200 μm (Wohlwend Engineering). For sponges, tiny pieces of *O. carmela* were excised and mixed with 20% BSA made up in seawater before transferring to 200 μm-deep high pressure freezing planchettes. Freezing of both the choanoflagellate and sponge samples was done in a Bal-Tec HPM-010 high pressure freezer (Bal-Tec AG).

#### Freeze Substitution

High pressure frozen cells stored in liquid nitrogen were transferred to cryovials containing 1.5 ml of fixative consisting of 1% osmium tetroxide plus 0.1% uranyl acetate in acetone at liquid nitrogen temperature (−195°C) and processed for freeze substitution according to the method of McDonald and Webb [47,48]. Briefly, the cryovials containing fixative and cells were transferred to a cooled metal block at −195°C; the cold block was put into an insulated container such that the vials were horizontally oriented, and shaken on an orbital shaker operating at 125 rpm. After 3 hours the block/cells had warmed to 20° C and were ready for resin infiltration.

#### Resin Infiltration and Embedding

Resin infiltration was accomplished according to the method of McDonald [48]. Briefly, cells were rinsed 3X in pure acetone and infiltrated with Epon-Araldite resin in increasing increments of 25% over 30 min plus 3 changes of pure resin at 10 min each. Cells were removed from the planchettes at the beginning of the infiltration series, and spun down at 6,000 X g for 1 min between solution changes. The cells in pure resin were placed in between 2 PTFE-coated microscope slides and polymerized over 2 h in an oven set to 100°C.

#### Serial Sectioning

Cells/tissues were cut out from the thin layer of polymerized resin and remounted on blank resin blocks for sectioning. Serial sections of varying thicknesses between 70 - 150 nm were cut on a Reichert-Jung Ultracut E microtome picked up on 1 × 2 mm slot grids covered with a 0.6% Formvar film. Sections were post-stained with 1% aqueous uranyl acetate for 7 min and lead citrate [49] for 4 min.

#### Imaging

Images of cells on serial sections were taken on an FEI Tecnai 12 electron camera.

### 3D Reconstruction & Analysis

ssTEM sections were imported as z-stacks into the Fiji [50] plugin TrakEM2 [51] and automatically aligned using default parameters, except for increasing steps per octave scale to 5 and reducing maximal alignment error to 50 px. Alignments were manually curated and adjusted if deemed unsatisfactory. Organelles and subcellular compartments were manually segmented and 3D reconstructed by automatically merging traced features along the z-axis. Meshes were then preliminarily smoothed in TrakEM2 and exported into the open-source 3D software Blender 2.77 [52]. Heavy smoothing of the cell body in TrakEM2 sacrifices fine structures associated with cellular projections or does not remove all distinct z-layers, which exist as reconstruction artefacts. Therefore, cell bodies were manually smoothed using the F Smooth Sculpt Tool in Blender of final distinct z-layers for presentation purposes only (Figure S3). All organelles were subjected to the same smoothing parameters across individual cells. All analysis was conducted using unsmoothed, unprocessed meshes. Organelle volumes were automatically quantified by the TrakEM2 software and enumerated in Blender 2.77 by separating meshes in their total loose parts.

The microvillar collar and flagellum were excluded from volumetric analysis as their total, representative length could not be imaged at this magnification. Cytosolic volume was calculated by subtracting total organelle volume from cell body volume, and is inclusive of cytosol, ribosomes and unresolved smaller structures excluded from 3D reconstruction. Endocytotic vacuoles were distinguished from food vacuoles by connection to the extracellular medium in ssTEM’s or by localisation to a cell protrusion. Cells in rosette colonies are numbered in order of their appearance along the image stack z-axis. Rosette colony diameters were calculated by measuring the largest distance of the z-axis midsection. Bridge length was measured in one dimension along the bridge midsection. Mean vesicle diameters were calculated from 20 measurements (or as many as possible if the vesicle type was rare) from single cells.

### Data Analysis

Univariate differences in the volume and number of subcellular structures between the two cell types were evaluated using Two-Sample t-tests. Shapiro-Wilk and Levene’s tests were used to assess normality and homogeneity of variance respectively. Statistical comparisons were conducted using data scaled against total cell volume. Correlations between colony cell number, cell volume and bridges per cell were assessed using Pearson correlation tests. All statistical analyses were conducted using R v 3.3.1 [53] implemented in RStudio v 0.99.903 [54].

